# Full-length mRNA sequencing reveals principles of poly(A) tail length control

**DOI:** 10.1101/547034

**Authors:** Ivano Legnini, Jonathan Alles, Nikos Karaiskos, Salah Ayoub, Nikolaus Rajewsky

**Affiliations:** Laboratory for Systems Biology of Gene Regulatory Elements, Berlin Institute for Medical Systems Biology (BIMSB), Max Delbrück Center for Molecular Medicine, Robert-Rössle Str 10, Berlin-Buch, Germany

## Abstract

Although mRNAs are key molecules for understanding life, there exists no method to determine the full-length sequence of endogenous mRNAs including their poly(A) tails. Moreover, although poly(A) tails can be modified in functionally important ways, there also exists no method to accurately sequence them. Here, we present FLAM-seq, a rapid and simple method for high-quality sequencing of entire mRNAs. We report a cDNA library preparation method coupled to single-molecule sequencing to perform FLAM-seq. Using human cell lines, brain organoids, and *C. elegans* we show that FLAM-seq delivers high-quality full-length mRNA sequences for thousands of different genes per sample. We find that (a) 3’ UTR length is correlated with poly(A) tail length, (b) alternative polyadenylation sites and alternative promoters for the same gene are linked to different tail lengths, (c) tails contain a significant number of cytosines. Thus, we provide a widely useful method and fundamental insights into poly(A) tail regulation.

## Introduction

Eukaryotic mRNA synthesis is a complex process that requires coordinated control of transcription, capping, intron splicing and 3’ end formation. For most mRNAs, 3’ end maturation involves the addition of a non-templated poly(A) tail, which plays a key role in nuclear export, translation initiation and turnover (reviewed in Nicholson et al., 2018, Jalkanen et al., 2014, Eckmann et al., 2010).

Poly(A) tails are thought to be synthesized by the rapid addition of ∼ 250 adenosines to the newly generated 3’ end of pre-mRNAs (Eckmann et al., 2010). After export, cytoplasmic control of tail length is thought to mainly consist of deadenylation, which precedes and typically controls mRNA degradation (Brown et al., 1998, Yamashita et al., 2005, Meyer et al., 2004, Chen et al., 2011), even though cytoplasmic polyadenylation plays a role in specific conditions and cell types (Jalkanen et al., 2014).

An important contribution to the understanding of tail length control and its consequence on mRNA fate came from the development of methods for poly(A) tail length profiling at the genome-wide level, including PAL-seq and TAIL-seq (Subtelny et al., 2014 and Chang et al., 2014). By combining short read sequencing with advanced biochemical or computational approaches, these methods provide deep coverage, but cannot sequence the entire tail and cannot unequivocally report the mRNA isoform to which the tail is attached. Moreover, while PAL-seq enables accurate estimation of a broad range of tail lengths, TAIL-seq is limited to a detection limit of ∼ 230 nt but can determine terminal modifications of poly(A) tails, which were discovered to control mRNA stability (Lim et al., 2014, Chang et al., 2014, Morgan et al., 2017, Lim et al., 2018, Chang et al., 2018).

A variety of studies that made use of these technologies reported that steady state tail length of most mRNAs is much shorter than 250 nt (Subtelny et al., 2014 and Chang et al., 2014, Lima et al., 2017), with a median between 50 and 100 nt for most mRNAs, confirming previous observations obtained for specific mRNAs, or with *in vitro*-reconstituted systems (e.g. Brown et al., 1998, Yamashita et al., 2005). Additionally, global poly(A) tail profiling reported poor or even negative correlation of tail length with expression, half-life and ribosome occupancy of mRNAs, with the notable exception of specific biological processes such as early embryogenesis (Subtelny et al., 2014, Lim et al., 2016, Eichhorn et al., 2016).

Here, we describe a new method for high-throughput sequencing of polyadenylated RNAs in their entirety, including the transcription start site, the splicing pattern, the 3’ end and the poly(A) tail for each sequenced molecule. We termed this technique **f**ull-**l**ength poly(**A**) and **m**RNA **seq**uencing, or FLAM-seq. We validated FLAM-seq in many different ways, including Northern blotting, sequencing of cDNA and RNA standards, Hire-PAT assays (Bazzini et al., 2012), and comparison to transcription start site databases. By applying our method to a variety of biological samples, we confirmed previous reports showing that steady state poly(A) tails are typically short and that tail length negatively or poorly correlates with gene expression, turnover and ribosome occupancy. By providing full-length mRNA sequence including the poly(A) tail, FLAM-seq allows to reconstruct dependencies between different levels of gene regulation - in particular promoter choice, alternative splicing, 3’ UTR choice, and polyA tail length. Therefore, we were able to study coupling of tail length control with 3’ UTR choice and to some extent with transcription start site choice. Moreover, we report that poly(A) tails contain internal non-A nucleotides, mostly cytosines.

## Results

### Sequencing mRNAs from head to toe with FLAM-seq

We developed FLAM-seq (Fig. 1A), that leverages a custom-made cDNA preparation method with single molecule real time sequencing for rapid and simple transcriptome-wide full-length mRNA and poly(A) tail length profiling. Poly(A)-selected RNA is enzymatically tailed with a short stretch of mixed guanosines and inosines, which serves as a priming site for an anchored oligonucleotide carrying a Unique Molecular Identifier (UMI) and a PCR handle. RNA is then reverse transcribed in combination with a chemically modified template switching oligo (iso-TSO, Kapteyn et al., 2010) that allows tagging the RNA 5’ ends with a second PCR handle. The resulting cDNA is amplified via PCR and subjected to long read sequencing with the PacBio Sequel System (Fig. 1A). Reads mapped to a few representative genes are shown in Fig. 1B and S1A.

**Fig. 1.**
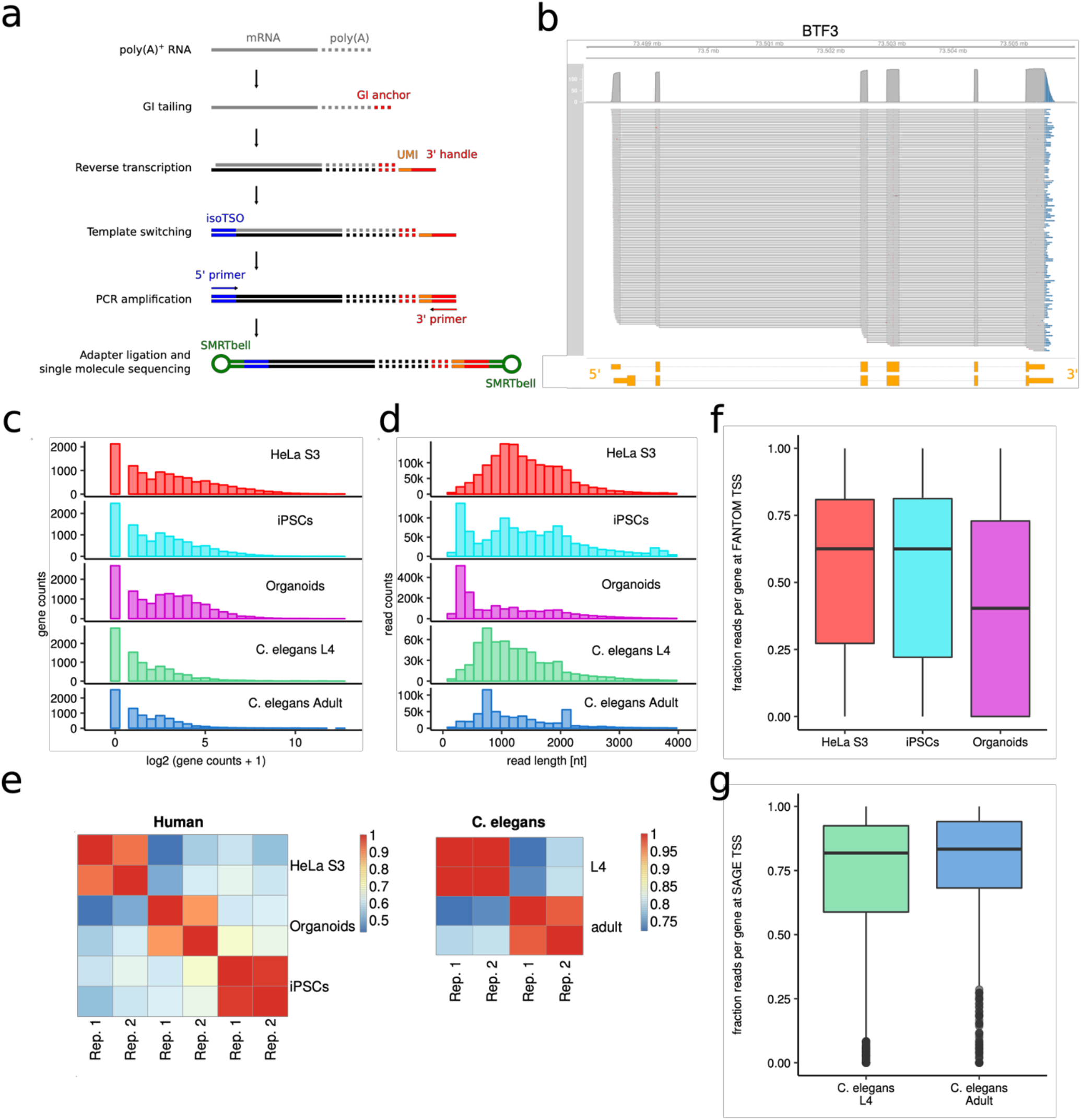
Full-length poly(A) mRNA sequencing (FLAM-seq). **A.** Outline of FLAM-seq. **B.** Genome-browser plot of aligned reads to a representative gene (BTF3). Non-templated poly(A) tail is shown at the 3’ end of the alignments in blue. Ensembl-annotated transcripts are shown at the bottom in yellow. **C.** Distribution of number of reads per gene for all sequenced samples (merged replicates). **D.** Distribution of FLAM-seq raw read length for all samples (merged replicates). **E.** Gene expression (reads per gene) correlation matrix between replicates of the human and *C. elegans* samples. **F.** Fraction of 5’ ends of reads overlapping with FANTOM5 transcription start sites (human) and **G.** SAGE TSS annotation data (*C. elegans*) per gene, for all samples (merged replicates).

To assess the outcome of FLAM-seq in a variety of biological systems, we sequenced two replicates of RNA from HeLa S3 cells, human induced pluripotent stem cells (iPSCs) and iPSCs-derived cerebral organoids on three PacBio SMRT-cells each, and two replicates of *C. elegans* L4 larvae and adult, egg-laying animals on two SMRT-cells each. For human and worm samples, we typically obtained sequences for circa 10000 and 7000 genes respectively, with a median read length of more than 1000 nt (Fig. 1C and D) and a good correlation of read counts per gene between the replicates (Fig. 1E). We compared the mapped 5’ ends of our reads with human transcription start sites (TSSs) annotated by the FANTOM5 project (Lizio et al., 2015) and observed that, on average, ∼ 40-70 % of the detected 5’ ends overlapped with them, depending on the sample (Fig. 1F). As a control, we trimmed the reads 5’ ends in silico and observed that the overlap with annotated TSSs was drastically reduced (Fig. S1B). Similar results were obtained with the worm samples, with ∼ 80% of the reads overlapping annotated TSSs (Fig. 1G and S1C). These data indicate that FLAM-seq is able to efficiently capture full-length RNAs. We note that coverage of longer transcripts is slightly biased towards their 3’ end, probably because of limitations in cDNA synthesis and PCR amplification (Fig. S1D).

### FLAM-seq accurately estimates poly(A) tail length

The advantage of FLAM-seq over existing full-length RNA sequencing technologies is that it captures the exact 3’ ends of the sequenced molecules. Since PacBio allows for sequencing of long homopolymers, we tested whether FLAM-seq could provide an accurate estimate of the length of the poly(A) tails. We generated a library from a pool of four synthetic cDNA standards, each carrying the PCR handles, a common sequence, a specific barcode and a poly(T) stretch of 30, 60, 90 and 120 nucleotides respectively (Fig. 2A). We sequenced this library and for each of the standards we observed a median tail length close to the expected one, with an offset of 2, 3, 5 and 6 nucleotides respectively and 97%, 92%, 75% and 78% of the reads having a tail within 10 nt from the expected length (Fig. 2A and B). To address possible artifacts occurring before the PCR amplification, we also produced one RNA standard by splint ligation of an *in vitro*-transcribed 200 nt-long RNA to a synthetic poly(A) stretch of 50 nt (Fig. 2A). When we sequenced the library generated from this RNA standard, we observed a median poly(A) length of exactly 50 nt and 78% of the tails within 10 nt from the median (Fig. 2A and B). To further assess the accuracy of our poly(A) length estimates, we first considered mitochondrial mRNAs, which have well-defined tails previously measured by other methods (Temperley et al., 2010), finding a substantial agreement between the known lengths and our HeLa datasets (Fig. S2A). Moreover, we performed a modified version of Hire-PAT (Bazzini et al., 2012), which consists of gene-specific PCR-amplification of the poly(A) tail followed by capillary electrophoresis, again observing substantial agreement with the sequencing data for five genes with different expression levels (Fig. 2C). We also assessed two of these genes by Northern blotting of RNA treated with RNase H together with a gene-specific oligo, with or without oligo-dT, in order to compare the electrophoretic mobility of the 3’ end of the transcript with and without the poly(A) tail. This PCR-free approach again confirmed the poly(A) length distributions observed by FLAM-seq (Fig. S2B).

**Fig. 2.**
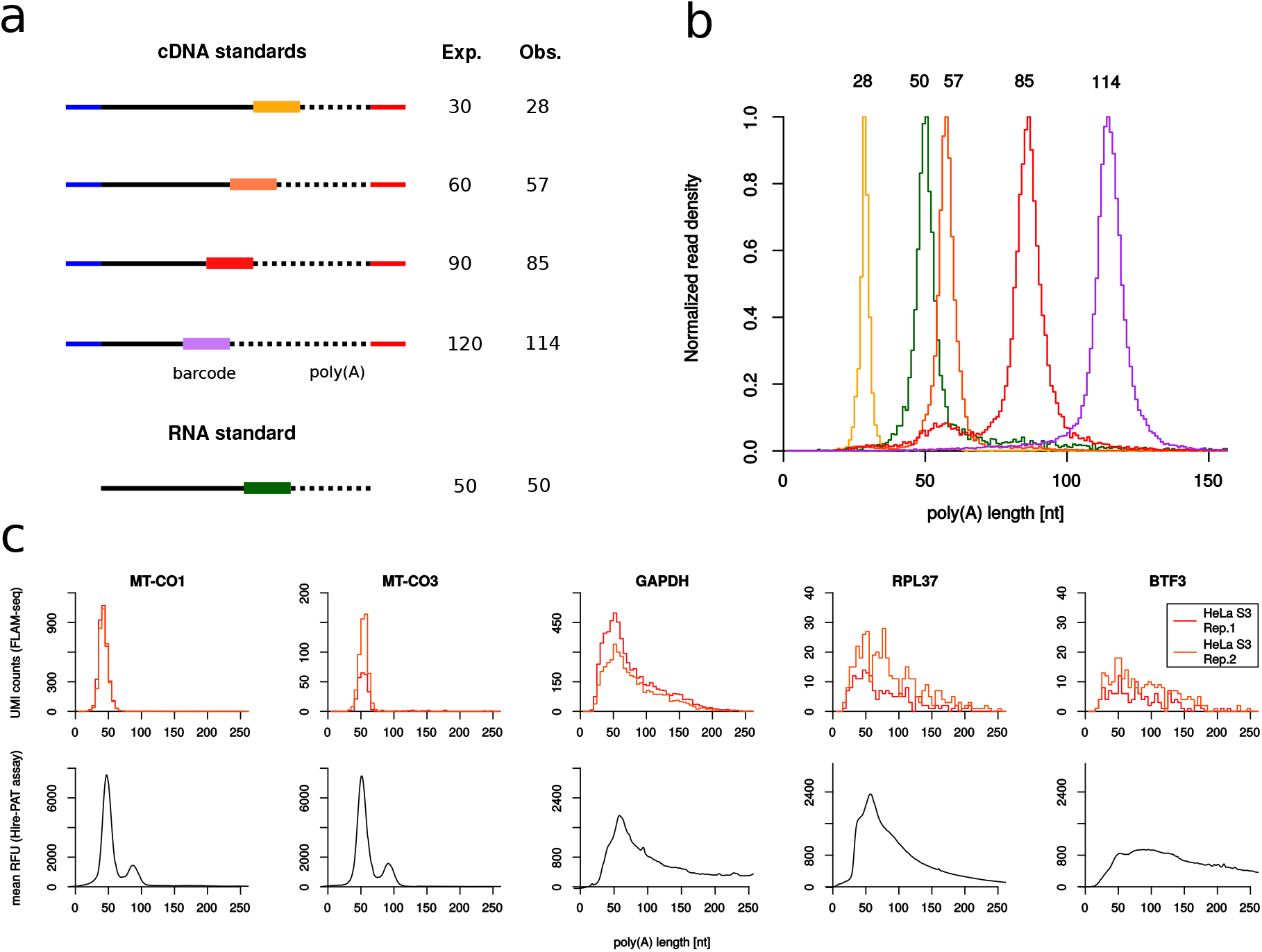
Validation of poly(A) length estimation by FLAM-seq. **A.** Sequence elements, including 10 nt barcodes, of four cDNA standards and one RNA standard with defined poly(A) length (Exp.). For each standard, the median poly(A) length observed with FLAM-seq is indicated (Obs.). **B.** Distribution of the measured poly(A) length for each standard. Line colors indicate each specific barcode as indicated in panel A. **C.** Poly(A) length validation by Hire-PAT assay for five genes in HeLa S3 cells. For each gene, the poly(A) length distribution measured with FLAM-seq in HeLa cells (two replicates in orange and red) are shown in the top panel and the PAT assay profile is shown in the bottom panel (mean of three replicates).

### Comparison of tail lengths across species, conditions and genomic features

Proven the accuracy of our method, we analyzed the poly(A) tail length profiles for each of the sequenced samples. The observed poly(A) length distributions per molecule (defined by UMIs) and per gene (defined by the length median of all UMIs attributed to said gene) were characteristic for each biological sample (Fig. 3A) and positively correlated among replicates (Fig. S3A).

**Fig. 3.**
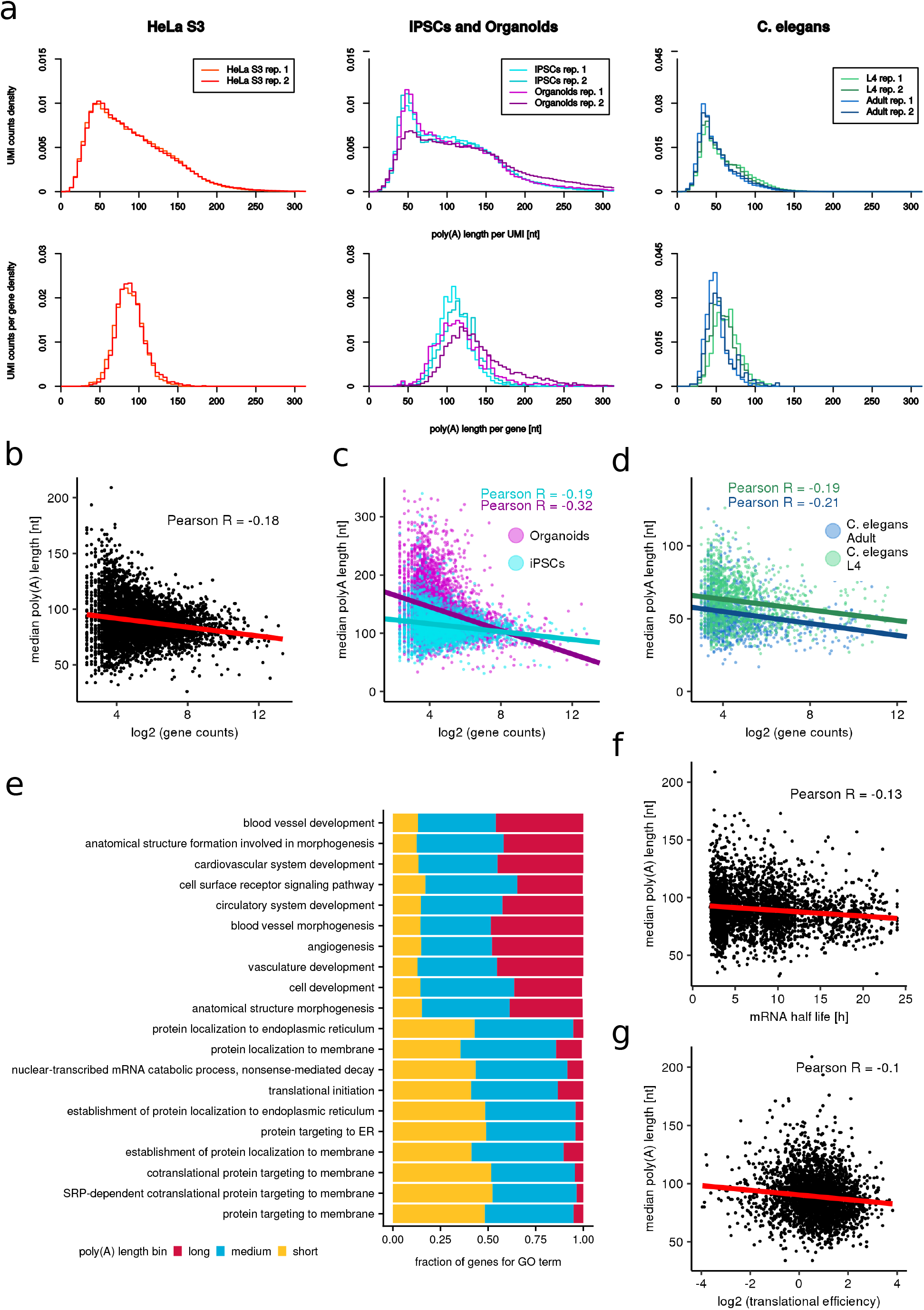
Poly(A) length profiles in human and *C. elegans* samples. **A.** Poly(A) length distribution across samples. Top panels: distribution per UMI, bottom panels: distribution per gene (median of UMIs for each gene). **B.** Correlation of median poly(A) tail length with expression level per gene (HeLa cells). **C.** in iPSCs and cerebral organoids and **D.** in *C. elegans* adult and L4. All described correlations are statistically significant (p < 0.001. Fisher Z-Transform/T-Test). **E.** GO terms enriched for genes binned by median Poly(A) length in Long, Medium and Short subsets. **F.** Correlation of median poly(A) tail length with mRNA half-life (Tani et al., 2012) per gene in HeLa cells (p < 0.001. Fisher Z-Transform/T-Test) **G.** Correlation of median poly(A) tail length with mRNA translational efficiency (Subtelny et al., 2014) per gene in HeLa cells (p < 0.001. Fisher Z-Transform/T-Test). A threshold of at least 10 UMIs per gene was applied in all analyses shown here.

Median poly(A) tail length per transcript (UMI) was between 81 and 114 nt for human samples, between 43 and 53 nt for worm samples, while median tail length per gene was slightly higher, between 86 and 131 nt for human samples and between 47 and 62 nt for worm samples (Fig. S3B). When comparing FLAM-seq data with other methods for poly(A) length estimation gene by gene, we observed low although positive correlation with PAL-seq and very poor correlation with TAIL-seq for HeLa cells, while good correlation with TAIL-seq for *C. elegans* (Fig. S3C and D). Perhaps surprisingly, also PAL-seq and TAIL-seq data poorly correlated between each other (Fig. S3E). Besides possible distinct technical biases in the three methods, as well as the different sequencing depths, these discrepancies might also be partly due to the diversity in the transcriptome of HeLa cells from various sources.

When comparing poly(A) tail lengths between iPS cells and organoids, we found 665 and 113 genes with statistically significant longer or shorter tails respectively. In *C. elegans* L4 and adult animals, we instead found 32 genes with longer tails and 136 genes with shorter tails. Interestingly, the differences in poly(A) tail length detected between iPSCs and organoids, as well as those detected in L4 and adult worms, were not correlated to changes in RNA abundance, indicating that poly(A) length control is not a major determinant of mRNA expression regulation in those systems (Fig. S3F, G, H and I).

Given the controversial association between gene expression, RNA turnover and translational efficiency with the poly(A) tail length, we sought to study the relationship between these different features. As also reported in a few recent studies (Lima et al., 2017, Subtelny et al., 2014), median poly(A) tail length negatively correlated with gene expression in all our datasets (Fig. 3B, C and D), with highly expressed genes typically having shorter tails. Along this line, Gene Ontology terms associated with transcripts having longer tails were enriched for regulatory functions such as developmental processes and signaling pathways, while shorter tails were associated with more housekeeping functions and especially with many terms related to membrane localization of proteins (Fig. 3E). Moreover, we observed a poor negative correlation of tail length with mRNA half-life and translational efficiency, suggesting that steady state poly(A) tail length does not seem to be a major determinant of mRNA stability or ribosome occupancy in the analyzed biological systems (Fig. 3F and G).

### FLAM-seq reveals poly(A) tail length dependency on 3’ UTR isoforms

One advantage of FLAM-seq is its capacity to sequence full-length transcripts, therefore allowing to study the relationship between the tails and other regulatory elements such as UTRs. We computed the tail length distributions for distinct mRNA isoforms, specifically for the ones having alternative polyadenylation sites and alternative transcription start sites. In human samples, we detected between 674 and 1005 genes with alternative 3’ UTRs, and among those between 290 and 547 having significantly different poly(A) length profiles between the detected isoforms (Fig. 4A). Surprisingly, we observed a statistically significant trend for isoforms deriving from distal APA site choice to carry longer poly(A) tails than proximal ones in the human samples (Fig. 4B, some examples in Fig. S4A). 3’ UTR length, in fact, positively correlates with median poly(A) length in all the analyzed samples (Fig. 4C, D and E). We also detected hundreds of genes with multiple TSS usage and amongst these a number of candidates where different TSS isoforms had statistically significant different poly(A) tail length profiles (Fig. 4F, two examples shown in Fig. S4B).

**Fig. 4.**
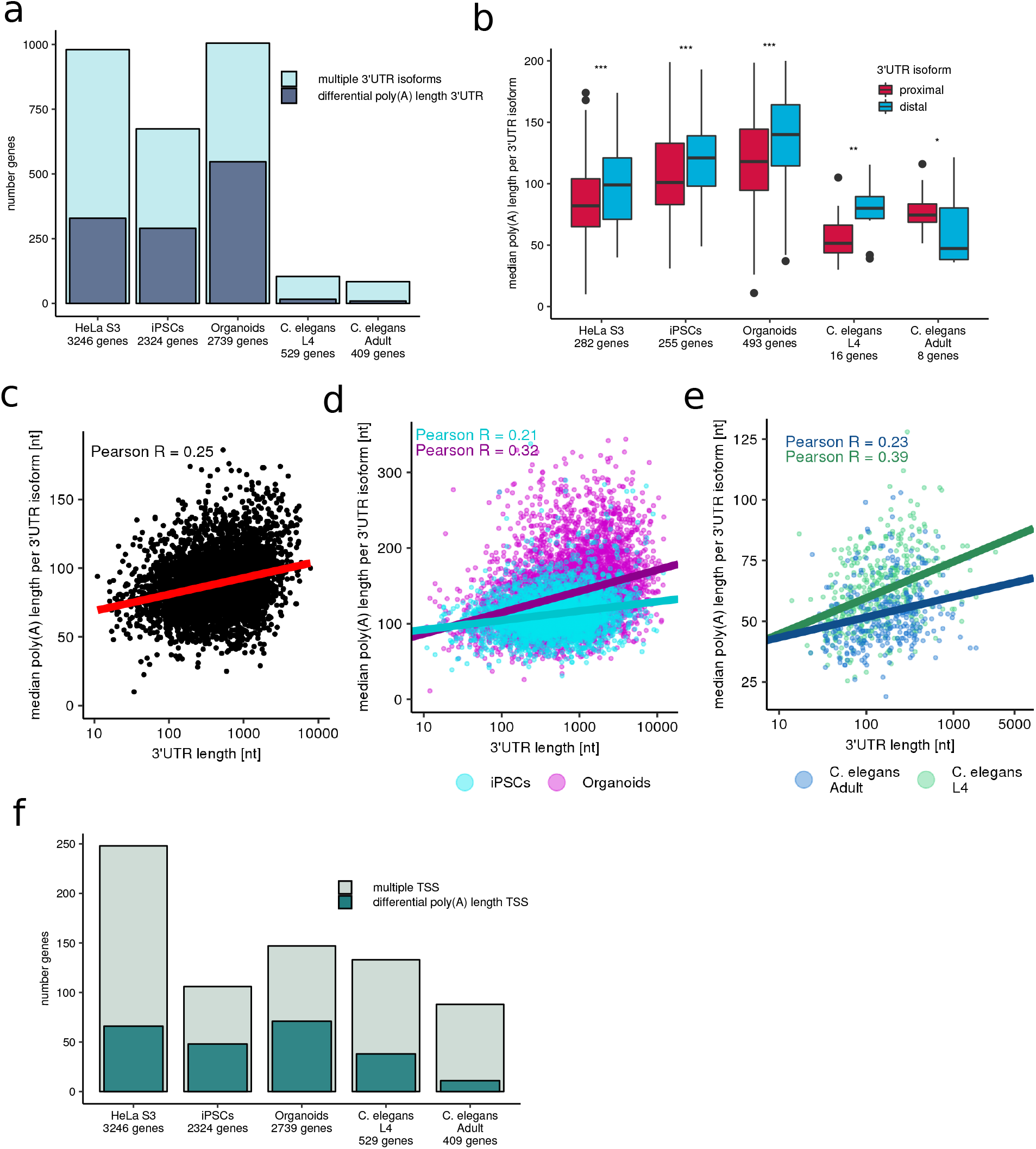
Widespread tail length dependency on mRNA isoform and internal nucleotide composition. **A.** Stacked bar plots represent the number of genes detected in each sample with more than one 3’ UTR isoform (light blue), and among those, genes with significantly different poly(A) tail length for different 3’ UTR isoforms (dark blue). The total gene sets considered for each sample are indicated below. **B**. Boxplots showing median poly(A) tail length per gene for proximal APA sites (red) and distal APA sites (blue) for all samples. (p values indicated; Wilcoxon-Test). **C.** Correlation of 3’ UTR length and median poly(A) length for all detected 3’ UTR isoforms in HeLa cells, **D.** iPSCs and cerebral organoids and **E.** *C. elegans* adult and L4 (p < 0.001; Fisher Z-Transform/T-Test). **F.** Stacked bar plots report the number of genes per sample with multiple Transcription Start Site (TSS) usage (light green) and the subset of genes with significant differences in poly(A) length distributions between UMIs associates with each TSS (dark green). Total gene numbers per sample are indicated below.

### Nucleotide composition of poly(A) tails

While PAL-seq and TAIL-seq can determine the size of poly(A) tails, FLAM-seq takes advantage of the PacBio technology, which enables to sequence long homopolymers. We analyzed the nucleotide composition of poly(A) tails and observed a fraction of non-A nucleotides of ∼ 0.3% and ∼ 0.2% in human and worm samples, which by far exceeded the observed frequencies of Cs and Us in the cDNA and RNA standards (Fig. 5A). Our data indicates the presence of such impurities in poly(A) tails, with cytosines being the most enriched non-A nucleotide in all samples.

**Fig. 5.**
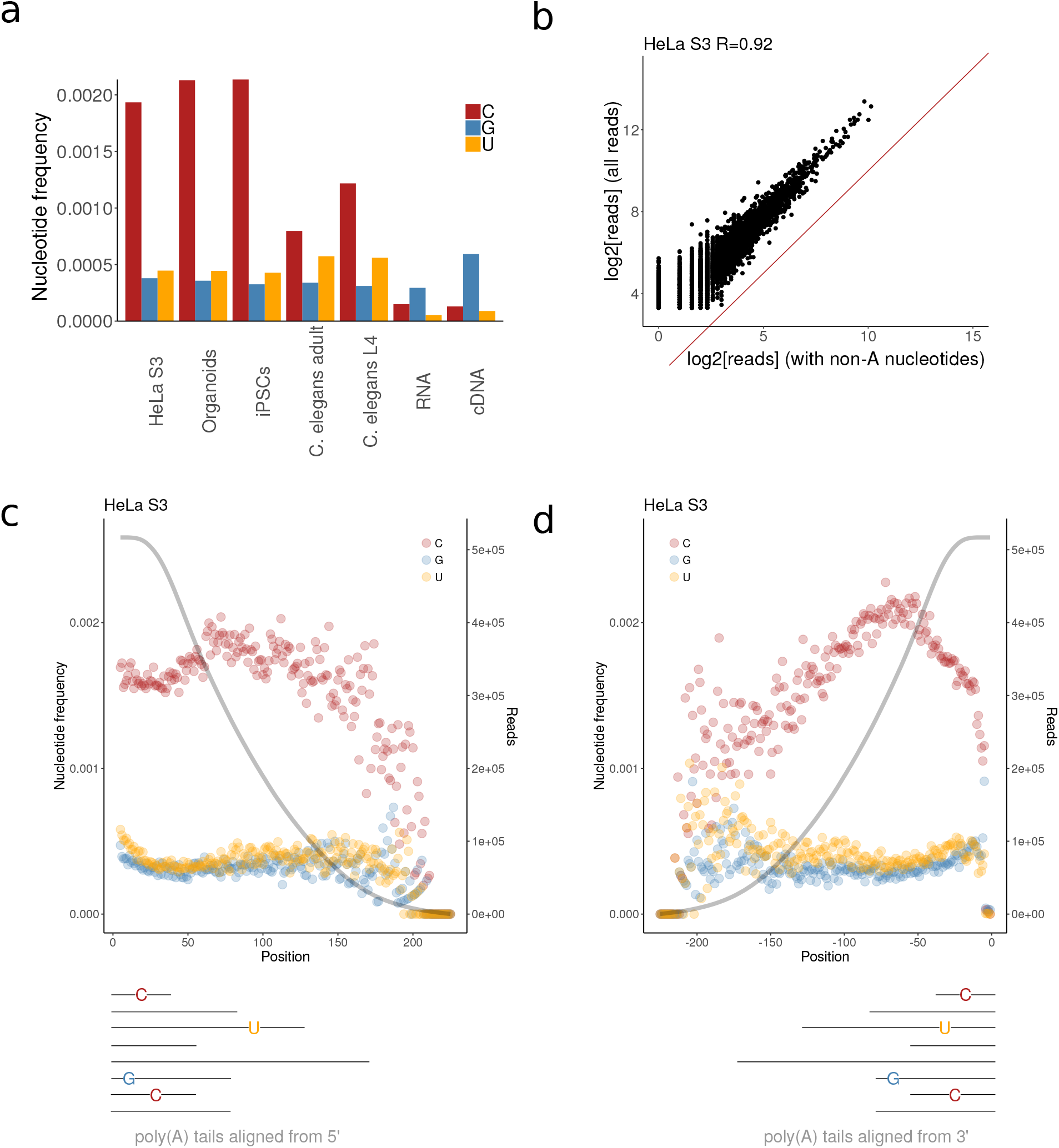
Nucleotide composition of poly(A) tails. **A.** Raw frequency of each non-A nucleotide within the poly(A) tails of the indicated samples. **B.** Correlation of number of reads containing impurities with total number of reads in HeLa cells. Red: Diagonal. **C.** Frequency of each non-A nucleotide for each position starting from the 10th nt of each tail, as sketched below and according to the color legend. In grey, total number of reads observed for each position. **D.** Frequency of each non-A nucleotide for each position starting from the last nt of each tail, as sketched below and according to the color legend. In grey, total number of reads observed for each position.

The number of non-A nucleotides for a given gene was highly correlated to the total number of molecules sequenced from that gene, indicating absence of gene-specificity (Fig. 5B and S5A).

In FLAM-seq, RNA is terminally tagged with a G/I tail, and the cDNA synthesis is primed with an oligonucleotide anchored with three Ts at the end of poly(A) tails. Therefore, we cannot interrogate our data for terminal modification of poly(A) tails, but we can observe more large-scale positional biases of non-A tail nucleotides. By aligning all tails either from their 5’ or 3’ end, we observed that cytosines generally dominate across all positions within tails but are slightly more enriched internally, with a depletion in the first or last 50 nt, especially in human samples (Fig. 5C and D, S5C). This effect seems significant when comparing it to tails in which the non-A nucleotides were randomly shuffled (Fig. S5B and C). Additional controls and observations are reported in the Supplementary Discussion.

## Discussion

The vast majority of eukaryotic mRNAs possess a 3’ poly(A) tail, which varies in length and composition. The fundamental importance of poly(A) tail length for gene activity has been reported and studied for decades. However, genome-wide analyses of tail lengths have been unavailable until recent years, when methods for genome-wide profiling of tails such as PAL-seq and TAIL-seq were published (Subtelny et al., and Chang et al., 2014). However, these methods have important limitations. Both methods provide only a short sequencing read (36 - 51 nt) for each mRNA to which the detected tail is attached. Thus, both methods do not provide information about the mRNAs beyond the tip of the 3’ UTRs to which the tails are attached, ruling out discovery of mRNA isoforms or analyses of how, for example, promoter choice is correlated to tail lengths. Moreover, PAL-seq has an extremely elaborate setup (in fact, we are not aware of any study that independently reproduced PAL-seq) and, by design, does not provide information about the tail sequence. TAIL-seq, however, is limited by the sequencing chemistry to study tails shorter than 250 nt, and provides useful sequence information only for the most terminal nucleotides of tails (10-30 nt), making it impossible to quantify the presence of internal non-A nucleotides in tails or to model their possible impact on deadenylation rates. For these reasons, we established FLAM-seq, a fast and simple full-length mRNA sequencing method that generates long reads with the PacBio Sequel system and provides information on mRNA isoform, poly(A) tail length and sequence for thousands of different transcripts. We applied FLAM-seq to a variety of biological samples and showed that it is able to accurately estimate the poly(A) tail length distribution of well-expressed genes. Besides global and accurate tail length profiling, which can be obtained with other technologies at higher coverage, FLAM-seq has two major advantages: a) it produces full-length mRNA sequences with an error rate lower than ∼ 0.8 % (a conservative estimate from mismatch and indel rates of reads mapped to the *C. elegans* genome) and b) it provides the sequence of poly(A) tails, with a baseline frequency of non-A nucleotides of ∼ 0.04 - 0.08%, as estimated from synthetic RNA and cDNA standards. Thanks to long read sequencing, mRNA isoforms can be easily discerned and poly(A) tails assigned to them without using reference gene models (e.g. as in Subtelny et al., 2014 and Lima et al., 2017). We focused on alternative polyadenylation (APA) sites, and observed hundreds of cases where APA choice led to changes in tail length for isoforms of the same gene. Moreover, we observed that distal APA sites are generally associated with longer tails than proximal ones and that 3’ UTR length positively correlates with tail length. Whether these changes reflect a difference in tail synthesis or result from different turnover and deadenylation kinetics remains to be established. The usage of alternative transcription start sites is thought to be generally uncoupled with 3’ end formation, but significant exceptions where promoter usage can influence APA choice can be found (e.g. Oktaba et al., 2015). In our data, transcription start site choice does not seem to influence APA usage, but we were able to assign poly(A) tails to mRNA isoforms produced from alternative TSS and observed a consistent fraction of these having a significant difference in tail length, showing that transcriptional and post-transcriptional regulatory elements in a gene body depend on each other.

Thanks to TAIL-seq, a number of terminal modifications of poly(A) tails were reported to play a role in mRNA decay (Lim et al., 2014, Chang et al., 2014, Morgan et al., 2017, Lim et al., 2018, Chang et al., 2018). However, TAIL-seq cannot be used to detect tail modifications within the vast majority of tail nucleotides as Illumina sequencing quality strongly deteriorates in homopolymeric stretches. With FLAM-seq, instead, we can sequence the entire tails (yet we cannot unambiguously identify 3’ terminal modifications). Comparison to our cDNA and RNA standards suggests that the detected non-A nucleotides are real and that poly(A) tails are not just stretches of adenosines. Cytosines were the most enriched, followed by uridines and guanosines. Notably, this ranking reflects the in-vitro incorporation rates of these nucleotides by the nuclear poly(A) polymerase (Wahle, 1991). The presence of non-A nucleotides within poly(A) tails can have substantial effects on tail shortening and therefore on mRNA turnover, as deadenylases preferentially trim adenosines (Lim et al., 2018). Since we did not observe any gene specificity, we did not find selective regulatory effects of tail impurities in the systems that we analyzed. However, these nucleotides in poly(A) tails might play significant roles in mRNA metabolism and require additional investigation (see supplementary discussion).

Very recently, Nanopore direct RNA sequencing has been used to estimate poly(A) tail length (Workman et al., bioRxiv 2018). Although its accuracy and precision were not yet thoroughly validated, and Nanopore sequencing has currently an extremely high error rate, it could in principle provide similar information to FLAM-seq, with the advantage of avoiding possible amplification biases but requiring significantly more input material and failing to provide accurate nucleotide identification within the tails. Moreover, we note that FLAM-seq libraries can also be sequenced with Nanopore devices (data not shown).

In conclusion, FLAM-seq provides for the first time global information on complete mRNA molecules at single nucleotide resolution. With its capacity of measuring global poly(A) tail profiles and full-length mRNA sequence, we suggest that it can be easily applied in combination with additional computational and experimental approaches to address relevant questions regarding the function and regulation of poly(A) tails and how, in general, transcriptional and post-transcriptional levels of gene regulation are coupled.

## Methods

### Cell culture

HeLa S3 cells were cultured in DMEM high glucose (cat. 41965, Thermo Fisher) supplemented with 10% FBS (cat. 10270106, Thermo Fisher).

iPSCs (XMO01 clone) were cultured in E8 flex medium (Thermo Fisher, cat. A2858501) and regularly passaged with Accutase (Stem Cell Technologies, cat. 07920). Cerebral organoids were generated according to the protocol described by Lancaster et al. (2014) with some modifications. Shortly, after dissociation into single-cell suspension with Accutase, 10000 cells were seeded per one well of 96 well plates in 100 µl of EB (medium supplemented with bFGF and 50 µM ROCK inhibitor). After 4 days, the medium was replaced by EB medium without bFGF and ROCK inhibitor and at day 6 by neural induction medium (NIM). At day 11, organoids were embedded in Matrigel (Corning, 356234) and kept in neural induction medium for two days, followed by organoid differentiation medium without retinoic acid (RA) for another four days. Next organoids were transferred to ultra-low attachment 6-well plates and culture on an orbital shaker (90 rpm) in organoid maturation medium. The composition of the original organoids maturation medium was changed by adding: chemically defined lipid concentrate (1x), ascorbic acid (0.4 mM), BDNF (20 ng/ml), HEPES.

### C. elegans

*Caenorhabditis elegans* animals (wild-type N2 Bristol) were cultured on OP50 bacteria, synchronized by bleaching, seeded and grown at 20 °C, collected in Trizol after 41 hrs (L4) or 84 hrs (adult) and homogenized using a Precellys 24 (Bertin Instruments).

### RNA preparation

All samples were collected in Trizol and then processed with Direct-zol RNA Miniprep kit (cat. R2052, Zymo research). Extraction and DNase treatment were performed according to the manufacturer protocol. Total RNA was checked on 2100 Bioanalyzer RNA picochip and then processed for poly(A) selection with the Truseq mRNA preparation kit (cat. RS-122-2102 Illumina), starting from 10 μg of RNA, adding 5 µl (1:100 ERCC Mix 1 cat: 4456740, Thermo Fisher). Poly(A)-selected RNA was again controlled on Bioanalyzer for excluding residual rRNA contaminations.

### Full-length mRNA library preparation

Poly(A)-selected RNA was tailed using the USB poly(A) length assay kit (cat. 764551KT, Thermo Fisher) in a 20 μl reaction with 14 μl RNA (equivalent to the initial 10 μg of total RNA), 4 μl tail buffer mix, 2 μl tail enzyme mix, for 1 hour at 37°C. Reaction was stopped with 1.5 μl of tail stop solution and 1 μl was checked on a picochip to exclude RNA degradation. Tailed RNA was then cleaned up with 1.8X RNAClean XP Beads (cat. A63987, Beckmann Coulter) and eluted in 18 μl H_2_O. For reverse transcription, RNA (16 μl) was incubated with 2 μl of one of the following RT primers:

GGTAATACGACTCACTATAGCGAGANNNNNNNNNNCCCCCCCCCTTT, TGAGTCGGCAGAGAACTGGCGAANNNNNNNNNNCCCCCCCCCTTT,

and then incubated for 3 minutes at 72°C, cooled down to 42 °C for 2 minutes and mixed with the following mix (SMARTScribe Reverse Transcriptase kit, cat. 639537, Clontech): 8 μl 5X First strand buffer, 1.5 μl DTT 20 mM, 4 μl dNTP mix 10 mM, 2 μl RNase inhibitor, 2 μl SMARTScribe RT, 2 μl H2O and 2 μl of the 12 μM iso-template switch oligo:

iCiGiCAAGCAGTGGTATCAACGCAGAGTACATrGrGrG.

The final mix was then incubated for 1 hour at 42°C and stopped for 10 minutes at 70°C. The resulting cDNA was purified with 0.6X XP DNA beads (cat. A63881, Beckmann Coulter) and resuspended in 42 μl of H_2_O. One μl of cDNA was checked on a picochip. The full-length mRNA library was then amplified by PCR with the Advantage 2 DNA polymerase mix (cat 639201, Clontech) according to the following mix: 10 μl 10X Advantage 2SA PCR buffer, 40 μl cDNA, 2 μl 10 mM dNTPs mix, 2 μl 5’ PCR primer IIA (from the kit), 2 μl 50X Advantage 2 Polymerase mix, 42 μl H_2_O, 2 μl of one of the two RT primer-matched universal reverse primers (10 μM):

GGTAATACGACTCACTATAGCGAG,

TGAGTCGGCAGAGAACTGGCGAA.

The PCR reaction was then performed in a thermocycler as follows: 98°C for 1 minute, 23 × (98°C for 10 seconds, 63°C for 15 seconds, 68°C for 3 minutes), 68°C for 10 minutes.

The amplified library was then purified with 0.6X Ampure XP DNA Beads and resuspended in 42 μl H_2_O. One μl of reaction was checked on fragment analyzer using High Sensitivity NGS Fragment Analysis Kit (cat. DNF-474, Advanced Analytical Technologies GmbH).

### PacBio sequencing

The purified PCR libraries were submitted to the Genomics core facility of MDC for PacBio sequencing. Sequencing libraries were prepared using the PacBio Amplicon Template Preparation and Sequencing Protocol (PN 100-081-600) and the SMRTbell Template Prep Kit 1.0-SPv3 according to the manufacturer’s guidelines. Sequencing on the Sequel was performed in Diffusion mode using the Sequel Binding and Internal Ctrl Kit 2.0. Every library was sequenced on 2 or 3 SMRT Cells 1M v2 with 1 × 600 min movie. Circular Consensus Sequence (CCS) reads were generated within the SMRT Link browser 5.0 (minimum full pass of 3 and minimum predicted accuracy of 90).

### RNA and cDNA standards

The RNA standard was produced by ligating a 200 nt *in vitro*-transcribed (IVT) RNA together with a synthetic poly(A) 50mer (custom-made by BioSynthesis). The IVT RNA was produced using pcDNA3.1(+) as template, amplified via PCR (10 × 100 μl reactions with GC buffer, Phusion High Fidelity DNA polymerase, cat. F530L, Life Technologies) with the following primers:

AAATAATACGACTCACTATAGGG,

AACGCTGGAACACGGGGGAGGGGCAAACAAC.

The PCR product was gel-purified and then used as template for in-vitro transcription in 10 × 20 μl reactions, each with 1 μg of DNA template, 1 μl 10 mM rNTPs, 2 μl 10X transcription buffer, 1 μl RNase inhibitor, 1 μl T7 RNA polymerase (from DIG RNA labeling kit, cat. 11175025910, Roche) in 20 μl total volume. Reactions were incubated for 2 hours at 37°C, then treated with 1 μl Turbo DNase (cat. AM2238, Thermo Fisher) for 15 minutes at 37°C and stopped for 5 minutes at 65°C. The resulting product was extracted with phenol-chloroform, ethanol-precipitated and gel-purified on 6% denaturing PAGE. The purified IVT RNA was then ligated with the poly(A) 50mer in 5 reactions each with 1 μg IVT RNA, 1 μg of RNA 50mer, 2 μl DNA splint oligo:

TTTTTTTTTTAACGCTGGAA,

which were incubated at 95°C for 2 minutes, slowly cooled to 25°C and mixed with 2 μl T4 DNA ligase (cat. M0202L, New England Biolabs), 2 μl DNA ligase Buffer and 0.5 μl RNase inhibitor. After 4 hours at 37°C, the reactions were pooled, purified with phenol-chloroform and precipitated with ethanol. Circa 10 μg of purified ligation were then used as input for library preparation as described before. The expected final sequence is pA_RNA_standard_50:

CAAGCAGTGGTATCAACGCAGAGTACATGGGAGACCCAAGCTGGCTAGCGTTTAAACTTAAGCTTGGTACCGAGCTCGGATCCACTAGTCCAGTGTGGTGGAATTCTGCAGATATCCAGCAGTGGCGGCCGCTCGAGTCTAGAGGGCCCGTTTAAACCCGCTGATCAGCCTCGACTGTGCCTTCTAGTTGCCAGCCATCTGTTGTTTGCCCCTCCCCCGTGTTCCAGCGTTGGGGGGGGGTCTCGCTATAGTGAGTCGTATTACCAAAAAAAAAAAAAAAAAAAAAAAAAAAAAAAAAAAAAAAAAAAAAAAAAA

cDNA standards were synthetized as ultramers by IDT:

pA_cDNA_standard_30: GGTAATACGACTCACTATAGCGAGACCCCCCCCCTTTTTTTTTTTTTTTTTTTTTTTTTTTTTTATGTCAGCAACACGGGGGAGGGGCAAACAACAGATGGCTGGCAACTAGAAGGCACAGTCGAGGCTGATCAGCGGGTTTAAACGGGCCCTCTAGACTCGAGCGGCCCCCATGTACT

CTGCGTTGATACCACTGCTTG

pA_cDNA_standard_60: GGTAATACGACTCACTATAGCGAGACCCCCCCCCTTTTTTTTTTTTTTTTTTTTTTTTTTTTTTTTTTTTTTTTTTTTTTTTTTTTTTTTTTTTAAGGTGCCACCACGGGGGAGGGGCAAACAACAGATGGCTGGCAACTAGAAGGCACAGTCGAGGCTGATCAGCGGGCCCATGTACTCTGCGTTGATACCACTGCTTG

pA_cDNA_standard_90: GGTAATACGACTCACTATAGCGAGACCCCCCCCCTTTTTTTTTTTTTTTTTTTTTTTTTTTTTTTTTTTTTTTTTTTTTTTTTTTTTTTTTTTTTTTTTTTTTTTTTTTTTTTTTTTTTTTTTTACTAGGTAGCCACGGGGGAGGGGCAAACAACAGATGGCTGGCAACCCCATGTACTCTGCGTTGATACCACTGCTTG

pA_cDNA_standard_120: GGTAATACGACTCACTATAGCGAGACCCCCCCCCTTTTTTTTTTTTTTTTTTTTTTTTTTTTTTTTTTTTTTTTTTTTTTTTTTTTTTTTTTTTTTTTTTTTTTTTTTTTTTTTTTTTTTTTTTTTTTTTTTTTTTTTTTTTTTTTTTTTTTTTAACAGTAGAACACGGCCCATGTACTCTGCGTTGATACCACTGCTTG

### Hire-PAT assays

Three replicates of HeLa S3 cells RNA were extracted, tailed and reverse-transcribed as described for the sequencing libraries. After the RT step, 1 μl of cDNA was used for each gene-specific PCR reaction, performed with Phusion High Fidelity DNA Polymerase and GC Buffer in 20 μl reactions (cat. F530L, Life Technologies) according to the manufacturer’s protocol as follows: 98°C for 30 seconds, 28 × (98°C for 10 seconds, 59°C for 15 seconds, 72°C for 3 minutes), 72°C for 2 minutes. In each reaction, the universal reverse primer was mixed together with one of the following gene-specific forward primers:

MT-CO1: ACCCCCCAAAGCTGGTTTC

MT-CO3: AAAAATAGATGTGGTTTGACTATTTC

GAPDH: AAAAAGCCTAGGGAGCCGCACCTTG

RPL37: CCAGTTCATCTTAAGAATGTCAACG

BTF3: GAAGAAGCCTGGGAATCAAGTTTG

After PCR, 1 μl of each reaction was then analyzed by capillary electrophoresis, with a Fragment Analyzer using High Sensitivity Small DNA Fragment Analysis Kit (cat: DNF-477, Advanced Analytical Technologies GmbH).

### Northern blots

For each Northern blot, three replicates of HeLa S3 cells RNA were extracted and treated as follows. Each RNA sample (20 μg) was split into two tubes and incubated with or without 5 μl of 1000 ng/μl oligo dT (15-18), 10 μl of 10 μM gene specific oligo, 6 μl of 5X RNase H buffer (250 mM Tris-HCl pH 7.5, 500 mM NaCl, and 50 mM MgCl_2_) and H_2_O to a final volume of 27 μl, incubated for 2 minutes at 85°C, 10 minutes at 42°C and then slowly cooled down to 32°C. To each mix, 1 μl Rnase inhibitor and 2 μl Hybridase(tm) Thermostable RNase H (cat. 108211, Biozym) were added and the reaction was incubated at 37°C for 1 hour. The treated RNA was then purified on 2X XP RNA beads, resuspended in 10 μl H_2_O, mixed with 1 volume of NorthernMax gly sample loading dye (cat. AM8551, Thermo Fisher), incubated for 30 minutes at 50°C and run on 2.5% 1X MOPS/0.7 % formaldehyde agarose gel (10X MOPS buffer: 0.2M MOPS, 50 mM Sodium Acetate, 10 mM EDTA, pH 7.0) in 1X MOPS buffer at 100 V for circa 2 hours. Each gel contained 3 replicates of Rnase H-treated samples, with or without oligo dT, and one non-treated lane. The gel was then imaged in a UV trans-illuminator for checking RNA integrity of the non-RNaseH-treated lane, soaked for 10 minutes in 0.5X TBE buffer, and blotted in a Biorad Trans-blot semidry system onto Hybond N+ membrane, between two layers of extra-thick paper previously soaked in 0.5X TBE. Blotting was performed for 45 minutes at 400 mA. After blotting, RNA was cross-linked with 265 nm UV at 2000 j/cm^2^ and the membrane was pre-hybdridized for 30 minutes at 58°C with Northernmax prehybridization/hybridization solution (cat. AM8677, Thermo Fisher) and hybridized overnight at the same temperature with a gene-specific DIG-labeled probe (100 ng/ml). The membrane was then washed twice in SSC 2X, 0.1 % SDS for 30 minutes at 58°C, twice in 0.2X SSC, 0.1 % SDS at 58°C for 30 and 60 minutes respectively. It was then blocked and hybridized with an anti-DIG AP-conjugated antibody (cat.11093274, Roche), washed and imaged with the CDP star ready-to-use system (cat. 12041677001, Roche), following the manufacturer’s protocol.

Gene specific Rnase H oligos were:

MT-CO3: CGCCTGATACTGGCATTTTG,

GAPDH: GTAGACCCCTTGAAGAGGGG.

Gene specific DIG-labeled probes were prepared by in-vitro transcription with the DIG RNA labeling kit (cat. 11277073910, Roche) according to the manufacturer’s protocol, on a PCR-derived DNA template obtained from HeLa S3 cDNA with the following primers:

MT-CO3_probe_fw: AAAAATAGATGTGGTTTGACTATTTC

MT-CO3_probe_t7_rv: TAATACGACTCACTATAGGGAAGACCCTCATCAATAGATG

GAPDH_probe_fw: AAAAAGCCTAGGGAGCCGCACCTTG

GAPDH_probe_t7_rv: TAATACGACTCACTATAGGGAGCACAGGGTACTTTATTG

## Data analysis

Raw reads were first filtered for keeping reads that have the characteristic subsequence [A]*10+[G]*9, or the reverse complementary sequence [C]*9+[T]*10, resulting from adding a GI-Tail to the Poly(A) Tail. Reads were oriented such that each read starts with the sequencing adapter sequence, followed by the UMI, the GI-Tail, the poly(A) tail and the sequence of the cDNA molecule. All reads correspond to cDNA orientation, i.e. they are reverse complement to RNA sequence of origin.

Poly(A) tail length for each read was determined by two different algorithmic approaches, applied with different parameters.

The first algorithm searches for at least n=10 subsequent T nucleotides with maximum 1 mismatch (A, C or G) and extends this putative poly(A) tail by iteratively searching for n+1 subsequent Ts. A maximum of n / T1 mismatches is allowed. T1 is a cutoff. After the maximum tail length for read is defined, the 3’ end of the putative poly(A) sequence is trimmed until a T nucleotide is reached.

The second algorithm utilizes a sliding window approach to identify the window where the relative frequency of T nucleotides drop below a threshold T2 within a sliding window of size L. Non-T nucleotides are clipped from the 3’ end of the remaining read until a TT dinucleotide is reached.

For each read, results from Algorithm 1 were obtained using parameters T1=25, T2=30, T1=35 and T1=40. Results from Algorithm 2 were obtained using parameter combinations L=20,T2=0.8; L=20,T2=0.85; L=25,T2=0.8; L=20,T2=0.8; L=20,T2=0.8 and L=20,T2=0.8

The consensus poly(A) tail sequences were determined by majority vote between the results produced by both algorithms and parameter combinations. The putative poly(A) tail sequence was removed from fastq reads, as well as the sequencing adapter sequences.

The remaining fraction of each read was mapped to human hg38, or *C. elegans* WB235 genome using STARLong (Dobin et al. 2013) with default parameters. (https://github.com/alexdobin/STAR/blob/master/bin/Linux_x86_64/STARlong)

Reads were assigned to Ensembl GRCh38.84 annotation gene models for human samples, or Ensembl WBcel235_82 gene models for *C. elegans* samples using FeatureCounts (Liao et al. 2014). For removal of reads resulting from internal priming of cDNA synthesis in A-rich transcript regions and sequences in the 5’ end of the putative poly(A) tails that are genomically encoded, raw fastq sequences were compared to the underlying genomic sequence.

The alignment indices of 100 nucleotides of the 3’ end of each alignment were mapped to positions in the raw fastq read. Raw fastq sequences were then from this position onwards compared to the underlying genomic sequence until a maximum of 3 subsequent mismatches is reached (thereby allowing for one Indel, as these occur frequently in PacBio data). Putative poly(A) tail sequences and length were then refined if necessary and alignment BAM files were further filtered by non-polyadenylated transcripts and/or transcripts resulting from internal priming. For downstream analysis, a minimum of 10 reads was required for each gene to be included in the analysis.

### Analysis of TSS and Relative Gene Coverage

Processed human CAGE TSS Peak files were downloaded (http://fantom.gsc.riken.jp/5/data/, Lizio et al., 2015) and converted to BED format. CAGE TSS and read alignments were sorted by gene model. For each alignment, the respective 5’-end was assigned to a CAGE TSS, if its 5’-start coordinate matched within a TSS peak boundary, or assigned to a ‘no_tss’ bin. For each gene, the fraction of annotated reads within TSS boundaries was computed relative to all reads for the gene.

For *C. elegans* samples, SAGE peak coordinates (Saito et al., 2013) were converted to BED format and peak coordinates within 10 nt windows were collapsed. *C. elegans* alignments were assigned as described above.

For control ‘in silico’ shortening, the respective 5’ alignment coordinate of read is subtracted by n with n ranging from 0 to 100 and relative fractions of reads within annotated TSS are computed for ‘truncated’ alignments as described above. Finally, for each value of n, median and standard deviation for relative TSS usage per gene are computed.

For metacoverage analysis, for each gene a ‘meta transcript’ was constructed by intersection of coordinates for all annotated exons. Coverage for each feature of this ‘meta transcript’ was computed using Bedtools Coverage (Quinlan et al. 2010). Coverage vectors for each feature of each ‘meta transcript’ were projected to a vector of uniform size for each gene. Genes with 0 coverage were removed. Each row (gene) in the resulting matrix was centered and scaled. Genes were binned by length and average metacoverage was computed for each bin.

### 3’UTR Annotation

For each alignment, the 3’ end coordinate with respect the RNA of origin was identified and sorted by gene. Putative 3’UTRs were predicted by peak detection on the alignment 3’-end coordinates specified for each gene using python peakutils module (https://bitbucket.org/lucashnegri/peakutils) using peakutils.indexes function with minimum peak height threshold parameter thres=0.1 and minimum peak distance parameter min_dist=30. Alignments were then sorted by predicted 3’ UTR ends. For identification of 3’ UTR start positions, splice sites for each read were first compared to annotated last exon starts for each gene, defined in human and *C. elegans* annotations. For identified last exons, 3’ UTR annotations within respective last exon coordinates used to defined 3’ UTR starts. 3’ UTR start for (truncated) alignment without splice sites were defined if unique 3’ UTR annotations were present for a gene. Corresponding Poly(A) Tail lengths of each alignment were mapped to each 3’ UTR isoform in order to infer isoform specific Poly(A) length distributions. For downstream analysis, a minimum of 5 reads per 3’ UTR isoform was required. Alternative 3’ UTR isoforms for each gene were compared by permutation testing, as described below in order to identify significant differences in Poly(A) length distributions.

### TSS Annotation

Alignments were assigned to human and *C. elegans* Transcription Start Sites as described above and poly(A) length distributions for each TSS were obtained by mapping poly(A) length to alignments assigned to each TSS. TSS with less than 5 reads assigned were discarded for further analysis. For each gene, the relative usage fraction of each TSS was computes as fraction of reads for the TSS compared to reads for all TSS. Multiple TSS usage for a gene was assumed if 2 or more TSS had a relative usage fraction of 0.2 - 0.8. TSS for genes with multiple TSS usage were compared for significantly differential Poly(A) length distribution by permutation testing as described below.

### Nucleotide composition of poly(A) tails

A small fraction of genes exhibited non-random di- or tri-nucleotide alterations proximal to the end of the 3’UTR. We identified those systematic alterations as 3’UTR remnants and therefore excluded the first 10 bases of each poly(A) tail for studying their nucleotide composition. The number of these cases was too small to influence the global poly(A) tail profiles, but had a non-negligible effect on the global nucleotide composition.

The positional frequencies of impurities were computed as follows. First, we counted the number of times a C, G or T was seen for a given position in all poly(A) tails of all genes. This number was divided by the number of times this position was ‘seen’ in the data, i.e. by the total number of poly(A) tails with length equal to, or greater than this position. This number was also plotted together with the positional frequencies to show our confidence in calling the latter ones (Fig. 5C and D and S5B and C).

To assess whether the observed impurity profiles reflect the biology of the tails, or are computational artifacts, we randomly shuffled the positional nucleotides of the poly(A) tails by keeping the length and the exact nucleotide composition of each tail intact.

### Normalization of consensus reads of cDNA and RNA standards

PacBio sequencing produces CCS reads by collapsing subreads generated along different iterations of the DNA polymerase. As the cDNA and RNA poly(A) standards have shorter lengths compared to sequenced molecules of biological samples, their consensus sequences would naturally have lower error rates because of higher number of subreads per CCS. In fact, the consensus sequences of molecules in the HeLa samples were produced by considering 6 times less individual reads, compared to the consensus sequences of the cDNA molecules. To assess whether this factor would be responsible for the nucleotide impurities observed in the poly(A) tails, we down-sampled the number of reads of the synthetic cDNA and RNA poly(A) standards to match the corresponding number of the HeLa samples. We generated the new CCS by running the SMRT link software on the down-sampled subreads with default parameters and quantified the composition of the poly(A) tails with our pipeline as before.

## Statistical Analysis

### Permutation Test differential Poly(A) Distributions

In order to test for differences in poly(A) tail length distributions between isoforms, genes or samples, a permutation test was computed on poly(A) tail lengths measured for individual molecules of each gene/isoform. A linear model was fitted on standard deviations of median poly(A) distributions for each cDNA standard of known length. For each measured poly(A) length L for a given molecule, the expected standard deviation inferred from the cDNA model length L is used to sample a new length value from a normal distribution with mean L. The median of the resampled distribution is then compared to the median inferred from the measured poly(A) tail length. A p-value is constructed from these comparisons by resampling the measured distributions 1000 times.

A minimum of 25 nt median difference between two Poly(A) distributions was required besides statistical significance for defining distributions as “different”.

## Author contributions

I.L., S.A. and J.A. conceived and optimized FLAM-seq. I.L., S.A. and J.A. performed all the experiments. J.A. and N.K. conceived and implemented the FLAM-seq analysis pipeline. J.A., N.K. and I.L. performed the computational analyses. N.R. contributed to both experimental and analyses design and supervised the whole project.

## Acknowledgements

We thank Agnieszka Rybak-Wolf for providing iPS cells and cerebral organoids samples, Jonathan Froehlich for providing *C. elegans* samples, Anastasiya Boltengagen for cell culture, Claudia Quedenau and Sascha Sauer of the BIMSB Genomics platform for adapter ligation, sequencing runs and preprocessing of sequencing data, Heiko Lickert for providing the XMO01 iPS cell line. We also thank all the members of the Nikolaus Rajewsky lab for critical and useful discussions. Ivano Legnini is recipient of an EMBO Long Term Fellowship (ALTF 1235-2016).

## Supplementary discussion

In search for non-A nucleotides within poly(A) tails, we analyzed the nucleotide content of tail sequences and observed the occurrence of Cs with higher frequency than other nucleotides, and above that observed in the RNA and cDNA standards.

Since our background model consists of synthetic molecules of defined lengths, we checked that poly(A) tails sequenced in HeLa cells with the same lengths within a window of +/- 5 nt retained the enrichments in non-A nucleotides (Fig. S5D).

A source of overestimation of impurities might be the consequence of the fact that our synthetic standards are shorter than the average cDNA amplicon from biological samples. In PacBio sequencing, each amplicon is circularized and sequenced multiple times, and the raw sequences (subreads) are then collapsed to give a high-confidence Circular Consensus Sequence (CCS). On average, shorter molecules can be sequenced more times, increasing the CCS accuracy. To test that if the impurities observed in biological samples are an artifact of different subread numbers, we sampled subreads of the cDNA and RNA standards to match the subread number distribution of the HeLa datasets and compared them again. Doing so, the frequency of cytosines increased but remained lower than those in the human and worm samples (Fig. S5E).

Finally, we report two observations regarding the relationship between C incorporation and tail length. First, when aligning the sequenced tails from either their 5’ or 3’ end, the frequency of Cs is increased in internal positions, especially after or before ∼ 50 nt from the end of the tails (Fig. 5C and S5B, C). Previous studies reported widespread terminal modifications of poly(A) tails with effects on RNA stability (Lim et al., 2014, Chang et al., 2014, Morgan et al., 2017, Lim et al., 2018, Chang et al., 2018). In the current setup of FLAM-seq, RNA is terminally tagged with a G/I tail and reverse transcription is performed with an oligo dC anchored to the tail with 3 Ts. Therefore, 3’ end modifications are excluded from our libraries, and guanylation cannot be distinguished from the artificially added G/I tails. However, non-G modifications can in principle be detected if reverse transcription is performed with a non-anchored oligo.

Second, we observed a positive correlation between tail length and frequency of Cs, implying a higher rate of incorporation for longer tails (Fig. S5F). When controlling for a similar pattern among the synthetic RNA and cDNA standards, we observed that the frequency of Cs is not uniform for different lengths of the same standard, possibly because of computational artifacts (Fig. S5G). However, the median frequency for each of the different standards does not seem to be dependent on their respective length, hinting at possible biological processes underlying the observed correlation (Fig. S5G).

---

**S1. Referred to Fig. 1.**
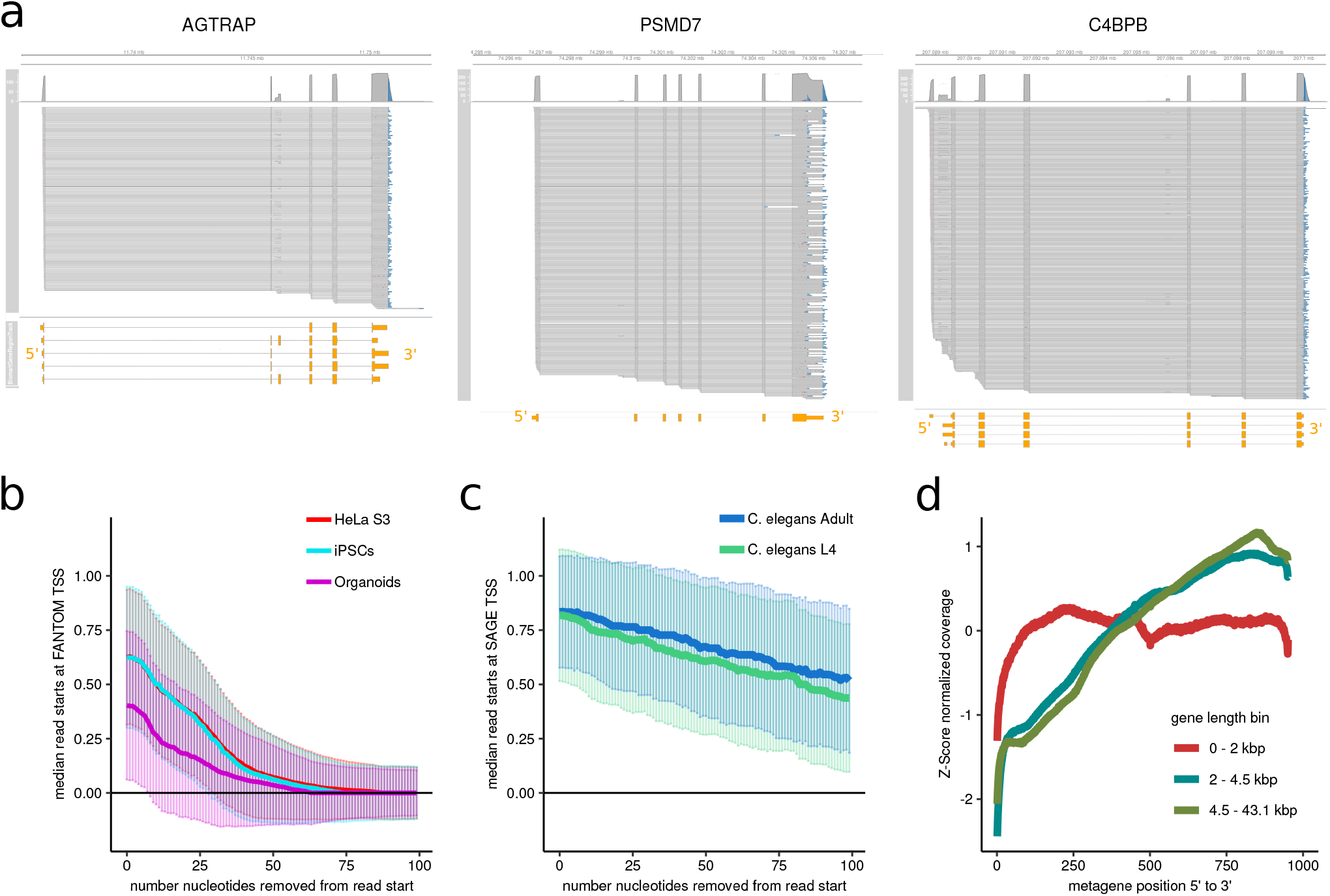
**A.** Genome browser-like plot of aligned reads for the indicated genes in HeLa cells, selected as paradigms of genes exhibiting alternative splicing, alternative termination or alternative transcription start site usage. Non-templated poly(A) tail is shown at the 3’ end of the alignments in blue. The Ensembl-annotated transcripts are shown at the bottom in yellow. **B**. Fraction of reads spanning the annotated TSS in human samples (FANTOM5 CAGE data) and **C.** *C. elegans* samples (SAGE data), after clipping the indicated number of nucleotides from the 5’ end of each alignment. **D.** Normalized meta-coverage of aligned reads from HeLa cells, binned by respective gene length.

**S2. Referred to Fig. 2.**
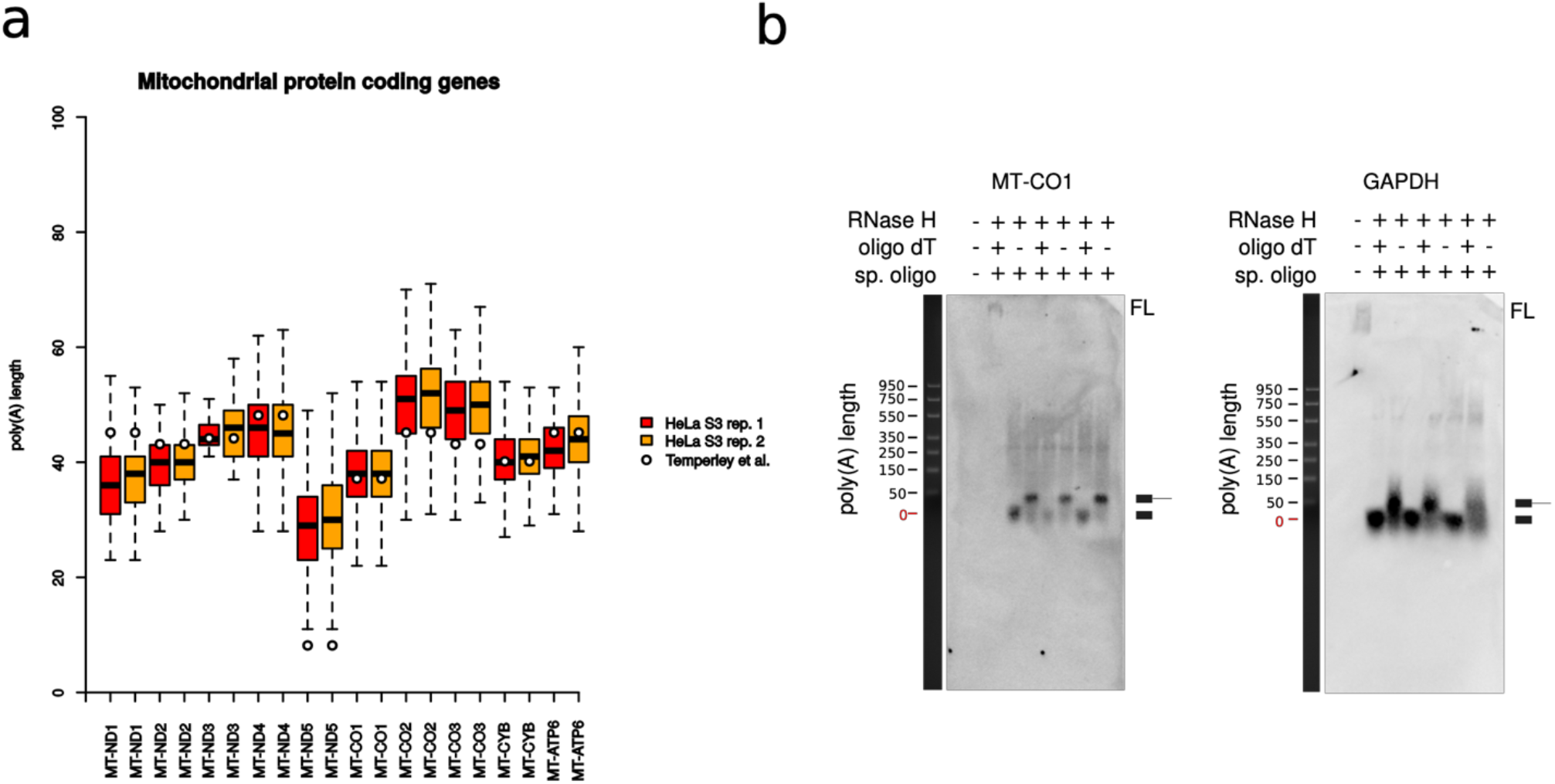
**A.** Boxplots of poly(A) tail length distributions for mitochondrial mRNAs, measured by FLAM-seq in the two HeLa replicates (orange and red, outliers not shown). Typical poly(A) length observed in HepG2 cells (Temperley et al., 2010) are plotted as white dots. Note: in Temperley et al., 2010, the MT-ND2 mRNA was reported to have a short poly(A) of 8 nt, which is below the threshold of 10 nt required in our analysis pipeline. The same mRNA was reported to exist in a second, less abundant form with a poly(A) tail between 25 and 50 nt, which is consistent with our estimate. **B.** Northern blot validation of poly(A) length for two genes in HeLa S3 cells. RNA is treated with RNase H in presence a gene specific oligo, with or without oligo-dT. On the left, the ladder is scaled according to the oligo-dT-treated samples to indicate the poly(A) length.

**S3. Referred to Fig. 3.**
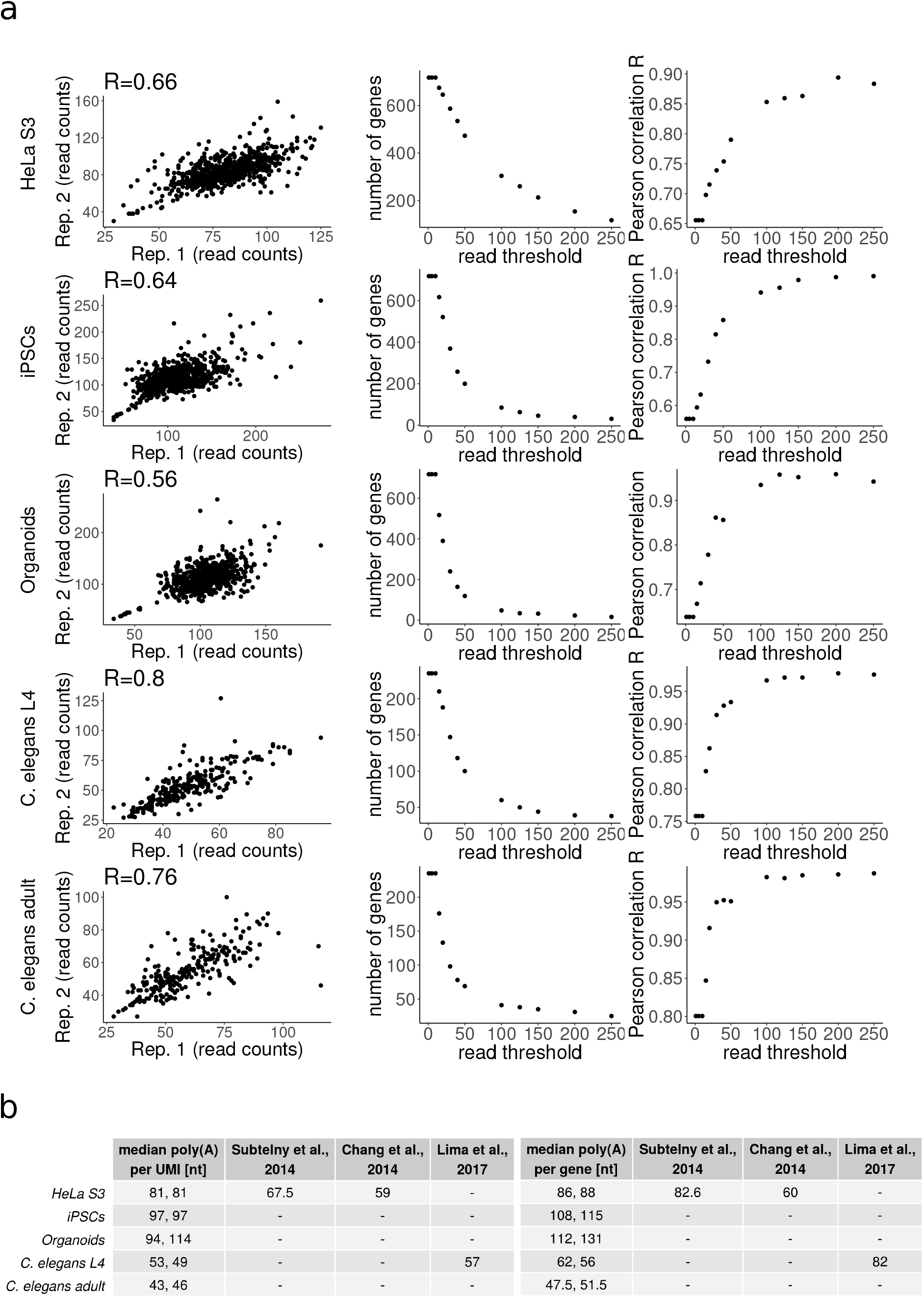
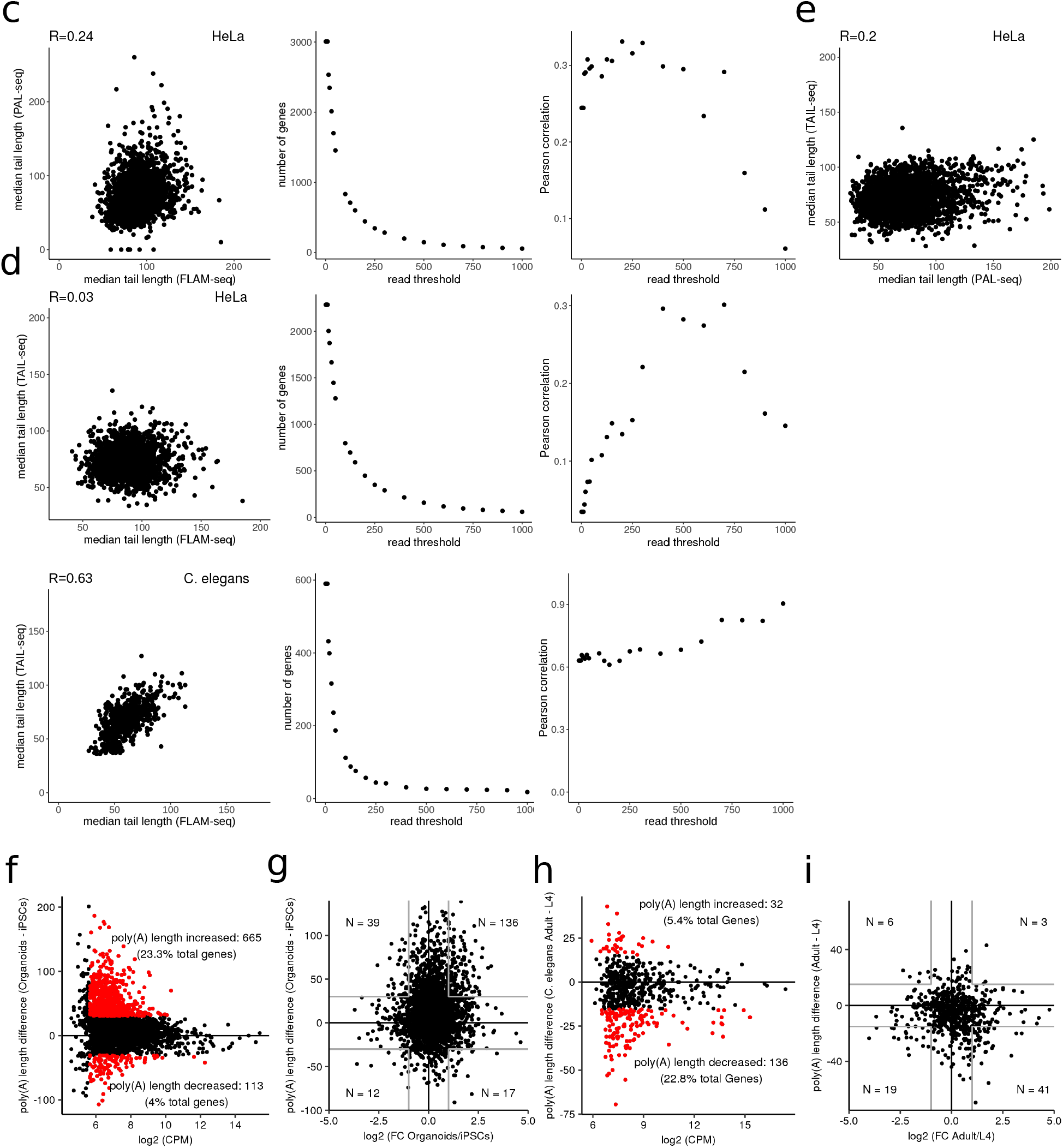
**A.** Left: correlation of poly(A) tail lengths between replicates for all samples in column, with a threshold of 10 UMIs/gene. On the side, the number of genes (center) and correlation coefficient (right) between poly(A) tail lengths between each couple of replicates for increasing thresholds of read counts per gene. **B.** Table showing median poly(A) tail length per UMI (left) and per gene (right) for FLAM-seq replicates in comparison with published data obtained with PAL-seq (Subtelny et al., 2014) and TAIL-seq (Chang et al., 2014 and Lima et al., 2017). **C.** Correlation of poly(A) tail lengths between FLAM-seq data and PAL-seq for HeLa cells (Subtelny et al., 2014), with a threshold of at least 10 UMIs/gene in FLAM-seq (left). On the side, number of genes (left) and correlation coefficient (right) between poly(A) tail lengths measured with FLAM-seq and PAL-seq for increasing thresholds of counts per gene. **D.** Top: same as C, for TAIL-seq data of HeLa cells (Chang et al., 2014). Bottom: same as C, for TAIL-seq data of *C. elegans* L4 stage (Lima et al., 2017). **E.** Correlation of poly(A) tail lengths between PAL-seq and TAIL-seq data for HeLa cells. **F.** Absolute Median Poly(A) length difference per gene by gene expression level between iPS and Organoid samples. In red genes with poly(A) tail differing for at least 30 nt between samples and with at least 10 counts per gene. **G.** Fold-change in gene expression level and median poly(A) length difference between genes between iPS and Organoids. **H.** and **I.** Same as F. and G. for *C. elegans* L4 and adult samples.

**S4. Referred to Fig. 4.**
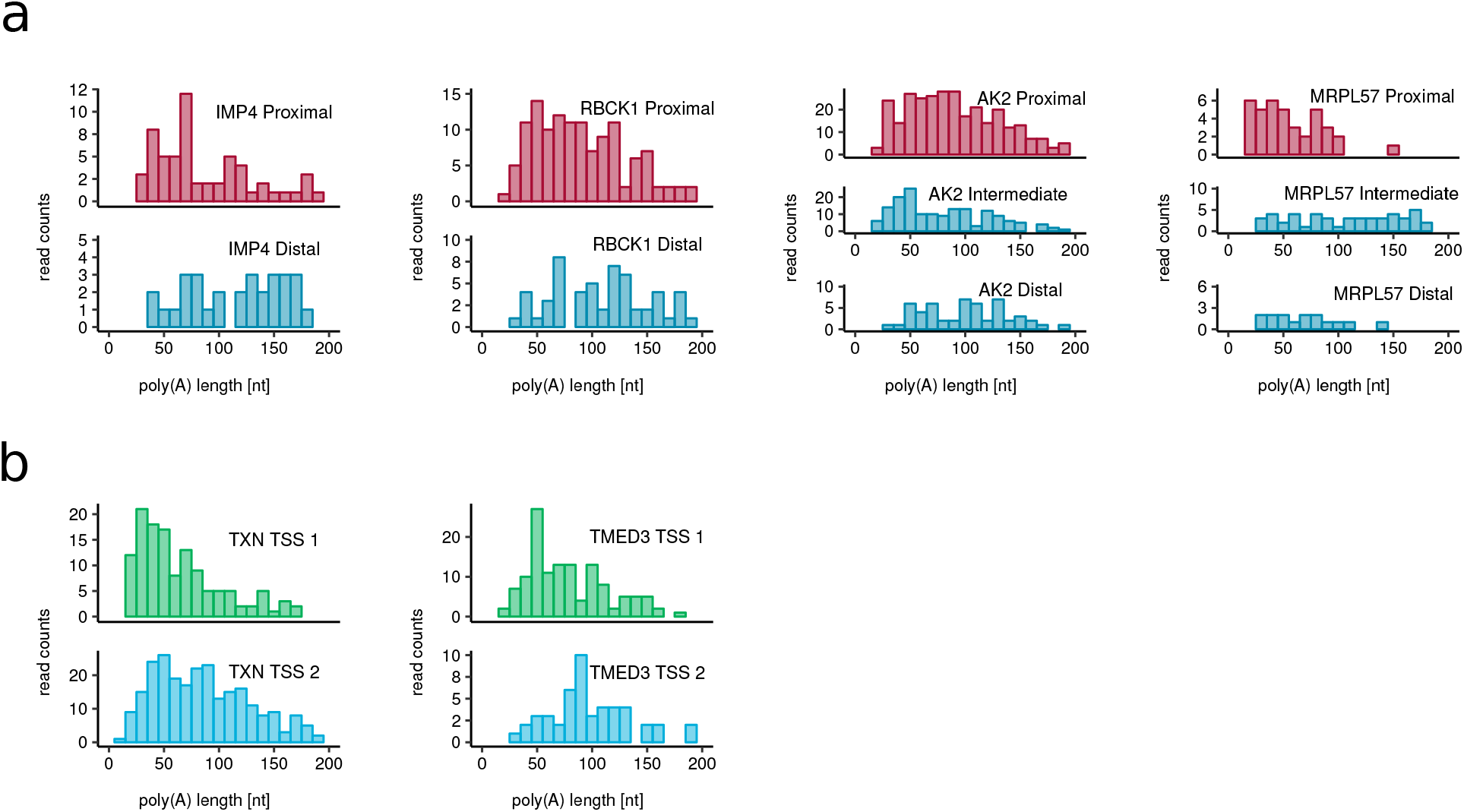
**A.** Poly(A) length distributions of four example genes having significantly different tail length between different 3’ UTR isoforms. **B.** Poly(A) length distributions of two example genes having significantly different tail length between different TSS isoforms. Statistical significance was assessed by the permutation test described in the methods section.

**S5. Referred to Fig. 5.**
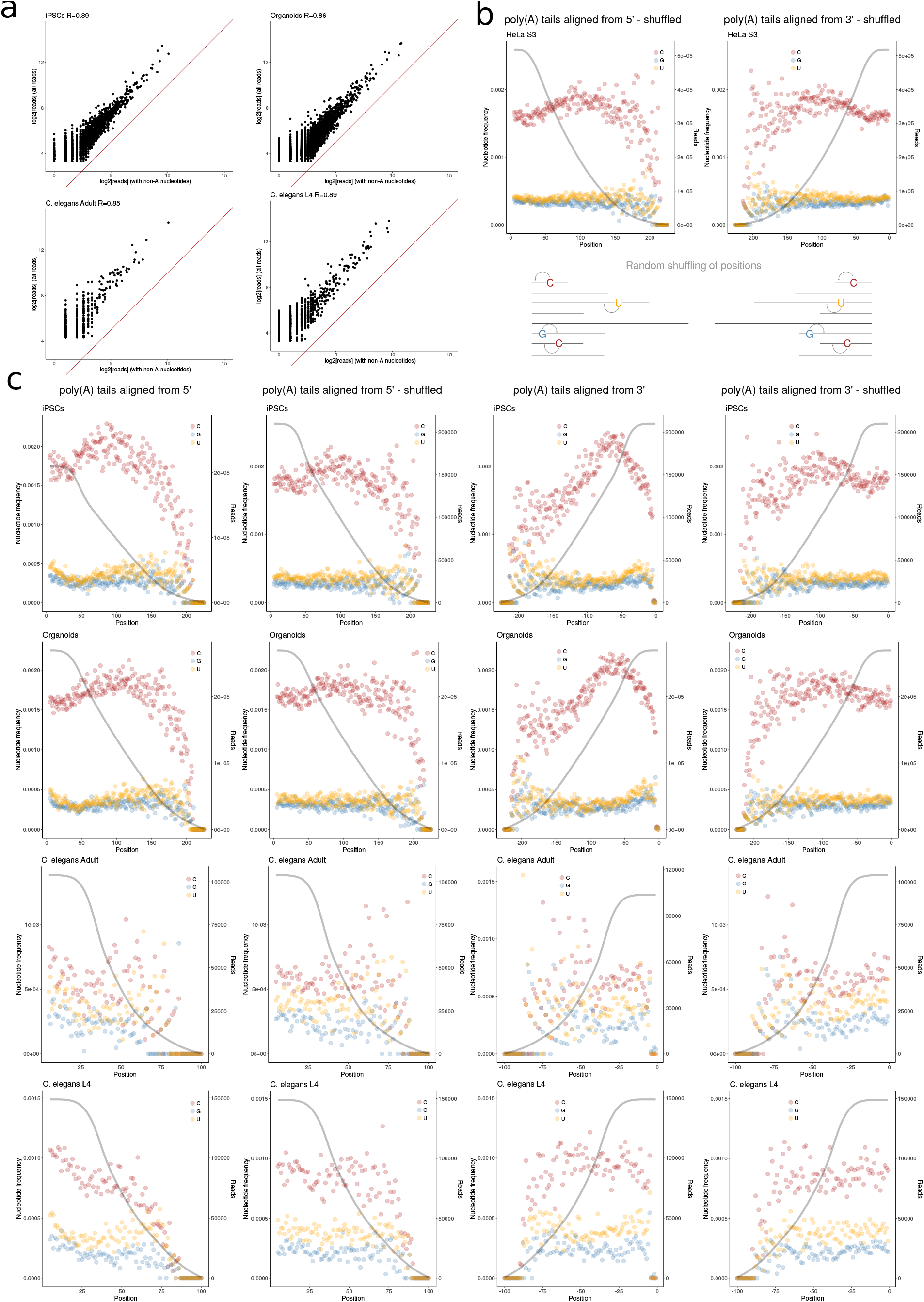
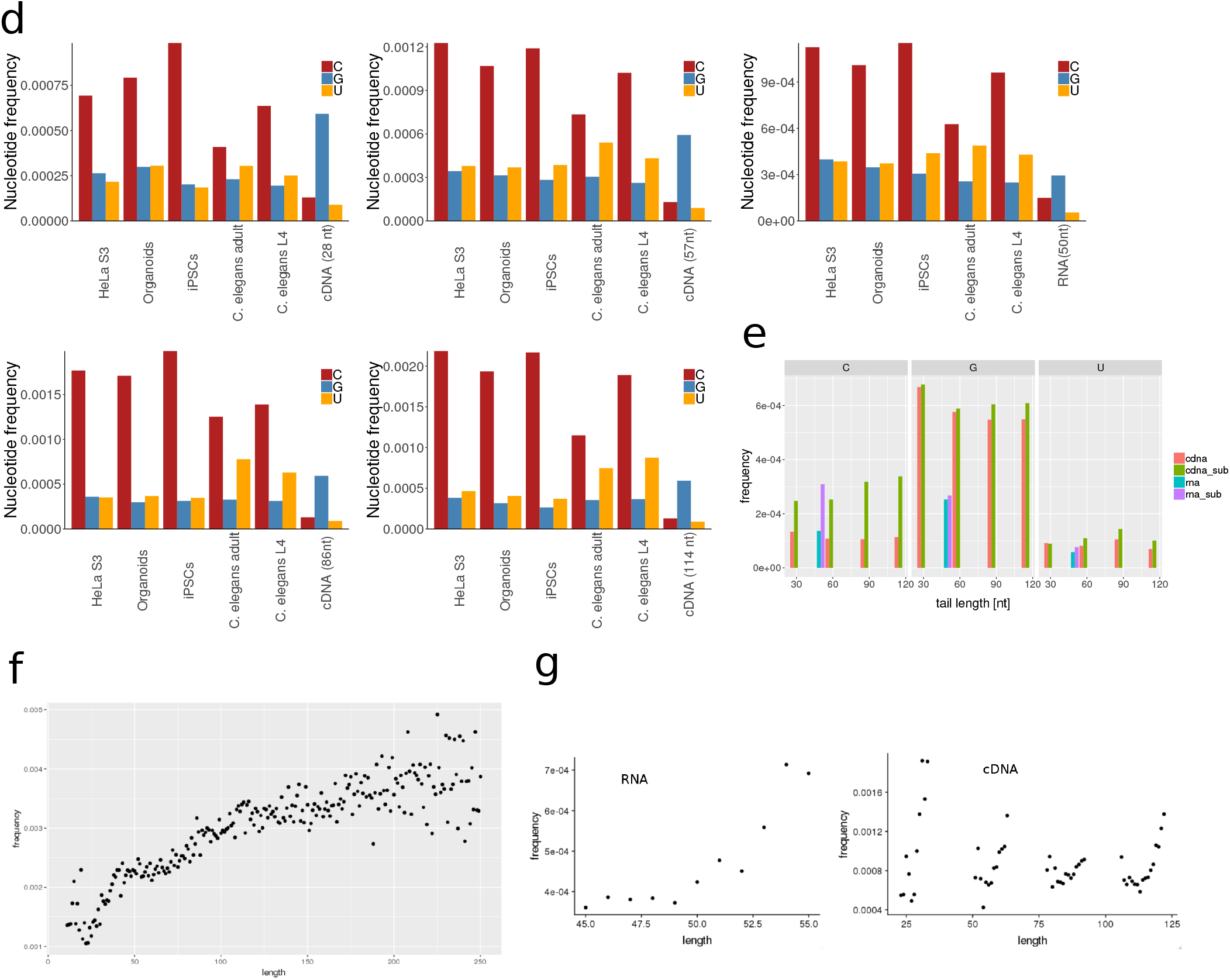
**A** Scatterplots showing correlation of reads containing modifications with all reads for iPSCs, organoids, *C. elegans* L4 and adult samples. **B.** Left: frequency of each non-A nucleotide is plotted for each position starting from the 10th nt of each tail, after randomly shuffling its position, as sketched below. Right: frequency of each non-A nucleotide is plotted for each position starting from the last nt of each tail, randomly shuffling its position as sketched below. In grey, total number of reads observed for each position. **C.** For the indicated samples, frequency of each non-A nucleotide for each position starting from the 10th nt of each tail (leftmost), frequency of each non-A nucleotide for each position starting from the 10th nt of each tail after randomly shuffling its position (center left), frequency of each non-A nucleotide for each position starting from the last nt of each tail (center right) and frequency of each non-A nucleotide for each position starting from the last nt of each tail after randomly shuffling its position (rightmost). In grey, total number of reads observed for each position. **D.** Bar plot showing the raw frequency of each non-A nucleotide within the poly(A) tails of the indicated samples, compared to each cDNA and RNA standard as indicated. For each plot, only tails of the sames length of the cDNA or RNA standard +/- 5 nt were taken into account. **E**. Frequency of each non-A nucleotide is plotted for cDNA and RNA standards of each length, as well as for cDNA and RNA standards after sub-sampling the same number of subreads as in the HeLa samples (cDNA sub and RNA sub). **F.** Scatterplot showing correlation of frequency of Cs with length of poly(A) tails in HeLa cells. **G.** Upper left panel: scatterplot showing correlation of frequency of Cs with tail length for the RNA standard, within 5 nt from the median observed length. Lower left panel: same as upper left, for all detected lengths. Upper right panel: Scatterplot for each of the cDNA standards showing correlation of frequency of Cs with tail length, within 5 nt from the median observed for each standard. Lower right panel: same as upper right, for all observed lengths.

